# Human Osteocytes Express MHC Class II and Act as Non-classical Antigen-Presenting Cells During Bacterial Infection

**DOI:** 10.64898/2026.08.28.747728

**Authors:** Mohammad Amjad Hossain, Dzenita Muratovic, Qi Sun, Boopalan Ramasamy, L. Bogdan Solomon, Plinio R. Hurtado, Gerald J. Atkins

## Abstract

Osteocytes are the most abundant cells in bone and are increasingly recognised not only for their role in skeletal remodelling and inflammatory signalling but also for their potential involvement in immune responses. In this study, we searched available gene expression datasets of human primary osteocyte-like cells exposed acutely to *Staphylococcus aureus* and identified significantly induced expression of key genes related to antigen processing and presentation. We then confirmed that human bone explant-derived osteoblastic cells, representative of a mature osteoblast-pre-osteocyte stage, expressed, as expected, high cell surface levels of major histocompatibility complex (MHC) Class I but also, low basal levels of the MHC Class II family member, HLA-DR. However, confocal imaging revealed high expression of MHC Class II molecules and the peptide-loading chaperone HLA-DM within the lysosomal compartments, consistent with canonical antigen-processing machinery. Differentiation towards a mature osteocyte phenotype increased MHC Class II protein levels and maintained expression of intracellular HLA-DM. Exposure of mature osteocyte-like cells to *S. aureus* further up-regulated both intracellular and cell surface MHC Class II expression. Demonstrative of antigen presenting cell functionality, *S. aureus*-exposed osteocytes induced autologous CD4^+^ T cell proliferation. Furthermore, MHC Class II expression in osteocytes was detected in bone sampled from patients with periprosthetic joint infections, providing evidence that these mechanisms operate *in vivo*. Together, our findings reveal that human osteocytes are capable of inducible MHC Class II-associated antigen presentation in response to bacterial challenge, pointing to a novel role for osteocytes in adaptive immune surveillance within bone.

**Lay Summary:** Osteocytes are the most common cell type in bone and are increasingly recognised as having important local and systemic regulatory functions. This study reveals a surprising new role: when challenged with bacteria, osteocytes can activate immune defence mechanisms normally associated with specialised immune cells. Specifically, human osteocytes express molecular signals that present to T cells, triggering a targeted immune response. We confirmed that this expression occurs in bone tissue taken from patients with infected joint replacements. These findings suggest that osteocytes play a key role in the immune response to bone infection, which may open new avenues for treating difficult-to-cure chronic bone infections.

## Introduction

Bone is increasingly recognised as an immunologically active tissue, in which skeletal cells interact dynamically with both innate and adaptive immune pathways, though this remains an emerging field.^(1)^ Osteoblasts and osteoclasts have been widely studied in the context of osteoimmunology,^(2,3)^ however, osteocytes remain comparatively underexplored despite comprising more than 90% of all bone cells. Embedded within the mineralised matrix and interconnected through the extensive lacunocanalicular network, osteocytes are ideally positioned to detect mechanical, metabolic and inflammatory signals and coordinate local skeletal responses.^(4–7)^ Traditionally regarded as mechanosensory regulators of bone remodelling, osteocytes are now understood to exert broader roles in mineral metabolism, endocrine signalling, and more recently, host defence. We recently reviewed how the osteoblast lineage, comprising osteoprogenitors, osteoblasts and osteocytes, functions as a stromal immune tissue capable of pathogen danger sensing, with osteoblasts in particular displaying inducible antigen-presenting cell (APC) capability.^(8)^

Osteocytes regulate bone turnover through secretion of key mediators, including sclerostin, RANKL, fibroblast growth factor-23 and various proinflammatory cytokines.^(7,9,10)^ Osteocyte differentiation is accompanied by distinct transcriptional and functional changes, marking the transition from matrix-forming osteoblasts to deeply embedded regulatory cells.^(11–15)^ Notably, despite being classically considered terminally differentiated, osteocytes can dedifferentiate into a proliferative and even migratory phenotype,^(13,16–19)^ likely reflecting their capacity to degrade their perilacunar matrix via osteocytic osteolysis/perilacunar remodelling.^(7,20,21)^

Osteocytes may also participate directly in host responses to bacterial infection.^(8,22–24)^ *Staphylococcus aureus*, the leading cause of osteomyelitis and periprosthetic joint infection (PJI), invades bone tissue, persists intracellularly, and evades antimicrobial therapy.^(25,26)^ Yang and Wijenayaka *et al.* demonstrated that osteocytes can harbour intracellular *S. aureus*, suggesting that these cells may act as a protected reservoir during chronic bone infection,^(23)^ a finding subsequently supported by other osteocyte-like cell models for *S. aureus* and additional pathogens.^(27–30)^

Consistent with this, evidence from PJI patients shows that osteocytes respond to infection in a distinct, pathogen-independent manner, triggering peri-lacunocanalicular matrix degradation and altered lacunar morphology, with lacunae becoming more circular and canalicular density significantly reduced.^(31,32)^ We recently showed in a mouse model of implant-related infection, that comparable changes occur within 12 days, correlating with the bacteria-to-host-cell ratio, rather than pathogen burden alone.^(33)^ Although the pathological significance of these changes remains unclear, they link pathogen sensing to localised tissue remodelling, suggesting an osteocyte-driven immune response and raising the question of whether osteocytes also engage in the adaptive arm of immunity.

Antigen presentation to CD4⁺ T lymphocytes is classically mediated by major histocompatibility complex Class II (MHC Class II) molecules, including human leukocyte antigen (HLA)-DR, together with intracellular peptide-loading regulators, such as HLA-DM.^(34,35)^ MHC Class II expression is typically restricted to professional APCs: dendritic cells,^(36)^ macrophages,^(37)^ and B cells.^(38)^ However, under inflammatory conditions, non-classical cell types including endothelial cells,^(39)^ fibroblasts,^(40)^ epithelial cells and osteoblast-lineage cells can acquire inducible MHC Class II expression.^(8,41,42)^ Osteoblasts exposed to inflammatory cytokines have indeed been shown to upregulate MHC Class II,^(43–46)^ suggesting that cells of skeletal lineage may contribute to local antigen presentation under pathological conditions. Whether this capacity extends to differentiated osteocytes has not been established.^(8)^

Given their abundance, longevity, strategic location, and capacity to sense microbial challenge, osteocytes are plausible but untested participants in local adaptive immune responses. This may be particularly relevant in chronic bone infection, where persistent bacterial antigens, inflammatory cytokines and immune cell infiltration create a microenvironment conducive to non-classical antigen presentation. We therefore aimed to determine whether human osteocytes express MHC Class II- and APC-associated pathways *in vitro* and *in vivo*, whether these responses are enhanced during osteocyte maturation or *S. aureus* exposure, and whether this expression is immunologically functional.

## Materials and Methods

### Gene expression database screening

Publicly available gene microarray datasets^(47)^ associated with our previous study,^(23)^ comparing human primary osteocytes acutely exposed to *S. aureus* with untreated controls, were manually screened for genes associated with antigen-processing and antigen-presentation. Fold-changes in infected relative to control, the direction of change and the false discovery rate -adjusted *p* values^(23)^ were recorded (**Table 1**).

**Table 1.**
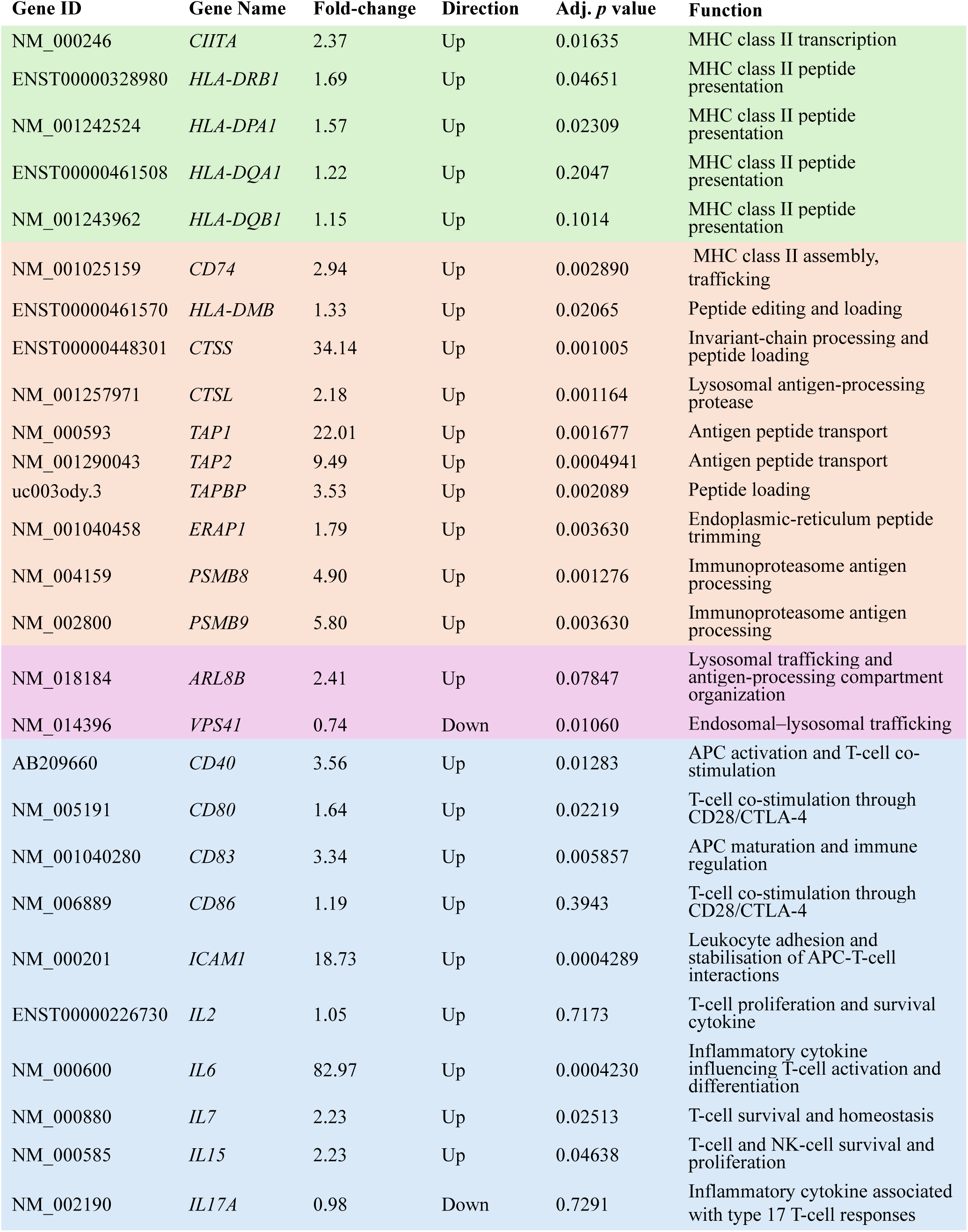
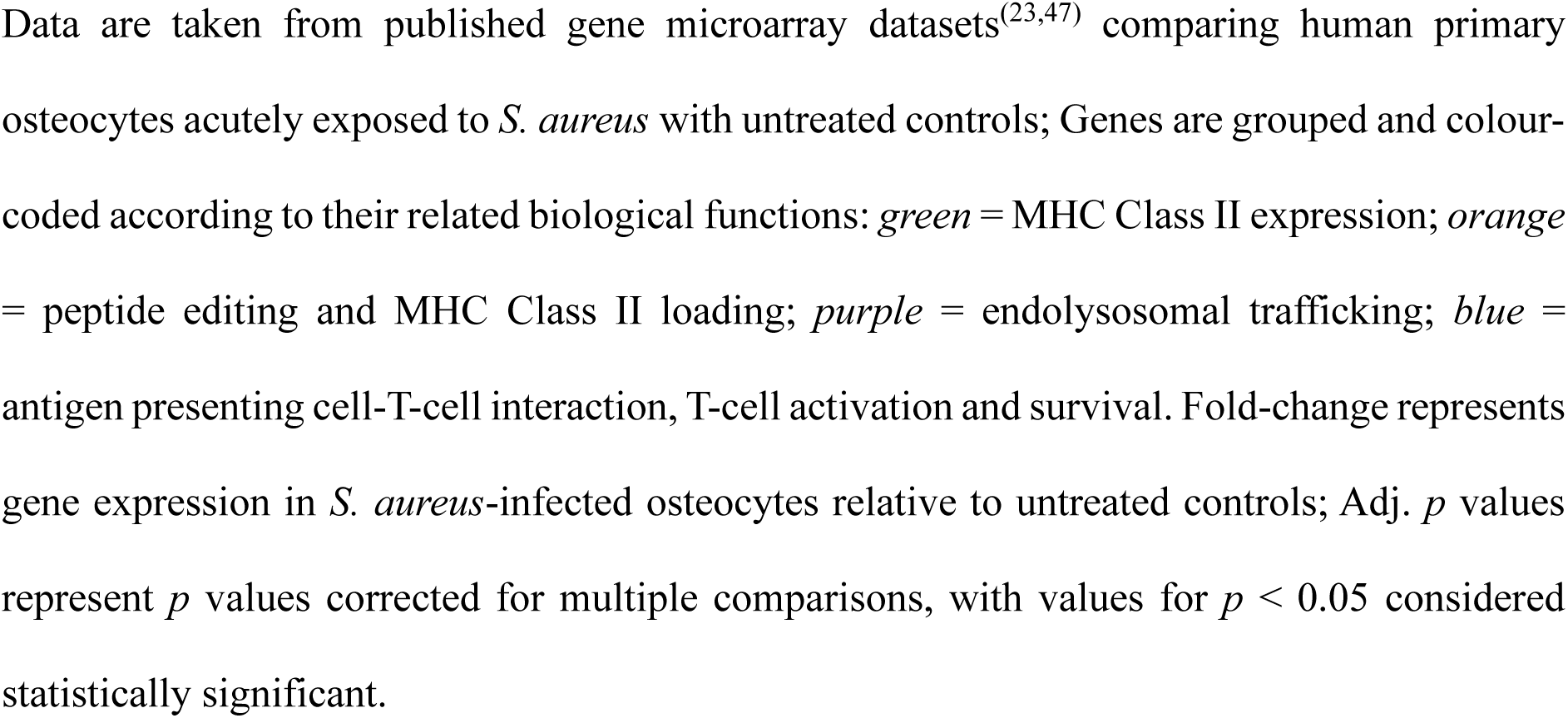
Microarray analysis of genes associated with antigen processing and presentation pathways in *S. aureus*-infected human osteocyte-like cells.

| Gene ID | Gene Name | Fold-change | Direction | Adj. *p* value | Function |
| --- | --- | --- | --- | --- | --- |
| NM_000246 | *CIITA* | 2.37 | Up | 0.01635 | MHC class II transcription |
| ENST00000328980 | *HLA-DRB1* | 1.69 | Up | 0.04651 | MHC class II peptide presentation |
| NM_001242524 | *HLA-DPA1* | 1.57 | Up | 0.02309 | MHC class II peptide presentation |
| ENST00000461508 | *HLA-DQA1* | 1.22 | Up | 0.2047 | MHC class II peptide presentation |
| NM_001243962 | *HLA-DQB1* | 1.15 | Up | 0.1014 | MHC class II peptide presentation |
| NM_001025159 | *CD74* | 2.94 | Up | 0.002890 | MHC class II assembly, trafficking |
| ENST00000461570 | *HLA-DMB* | 1.33 | Up | 0.02065 | Peptide editing and loading |
| ENST00000448301 | *CTSS* | 34.14 | Up | 0.001005 | Invariant-chain processing and peptide loading |
| NM_001257971 | *CTSL* | 2.18 | Up | 0.001164 | Lysosomal antigen-processing protease |
| NM_000593 | *TAP1* | 22.01 | Up | 0.001677 | Antigen peptide transport |
| NM_001290043 | *TAP2* | 9.49 | Up | 0.0004941 | Antigen peptide transport |
| uc003ody.3 | *TAPBP* | 3.53 | Up | 0.002089 | Peptide loading |
| NM_001040458 | *ERAP1* | 1.79 | Up | 0.003630 | Endoplasmic-reticulum peptide trimming |
| NM_004159 | *PSMB8* | 4.90 | Up | 0.001276 | Immunoproteasome antigen processing |
| NM_002800 | *PSMB9* | 5.80 | Up | 0.003630 | Immunoproteasome antigen processing |
| NM_018184 | *ARL8B* | 2.41 | Up | 0.07847 | Lysosomal trafficking and antigen-processing compartment organization |
| NM_014396 | *VPS41* | 0.74 | Down | 0.01060 | Endosomal–lysosomal trafficking |
| AB209660 | *CD40* | 3.56 | Up | 0.01283 | APC activation and T-cell co-stimulation |
| NM_005191 | *CD80* | 1.64 | Up | 0.02219 | T-cell co-stimulation through CD28/CTLA-4 |
| NM_001040280 | *CD83* | 3.34 | Up | 0.005857 | APC maturation and immune regulation |
| NM_006889 | *CD86* | 1.19 | Up | 0.3943 | T-cell co-stimulation through CD28/CTLA-4 |
| NM_000201 | *ICAM1* | 18.73 | Up | 0.0004289 | Leukocyte adhesion and stabilisation of APC-T-cell interactions |
| ENST00000226730 | *IL2* | 1.05 | Up | 0.7173 | T-cell proliferation and survival cytokine |
| NM_000600 | *IL6* | 82.97 | Up | 0.0004230 | Inflammatory cytokine influencing T-cell activation and differentiation |
| NM_000880 | *IL7* | 2.23 | Up | 0.02513 | T-cell survival and homeostasis |
| NM_000585 | *IL15* | 2.23 | Up | 0.04638 | T-cell and NK-cell survival and proliferation |
| NM_002190 | *IL17A* | 0.98 | Down | 0.7291 | Inflammatory cytokine associated with type 17 T-cell responses |

Data are taken from published gene microarray datasets<sup>(23,47)</sup> comparing human primary osteocytes acutely exposed to *S. aureus* with untreated controls; Genes are grouped and colour- coded according to their related biological functions: *green* = MHC Class II expression; *orange* = peptide editing and MHC Class II loading; *purple* = endolysosomal trafficking; *blue* = antigen presenting cell-T-cell interaction, T-cell activation and survival. Fold-change represents gene expression in *S. aureus*-infected osteocytes relative to untreated controls; Adj. *p* values represent *p* values corrected for multiple comparisons, with values for $p < 0.05$ considered statistically significant.

### Cell isolation and culture

All procedures involving the collection and use of human bone biopsy specimens blood samples, as well as associated clinical isolates, were approved by the Human Research Ethics Committee of the Royal Adelaide Hospital (Approval No. 14446).

Osteoblast-like normal human bone-derived cells (NHBC) were isolated from trabecular bone tissue as previously described.^(15,31)^ Bone specimens were obtained from patients undergoing total hip arthroplasty from Royal Adelaide Hospital, Australia. Trabecular bone was extensively washed with sterile phosphate-buffered saline (PBS) to remove residual blood and marrow components, then sectioned into small fragments (∼1–2 mm^3^) under sterile conditions. Bone chips were transferred to tissue culture flasks for outgrowth of NHBC.

The cells were maintained in growth medium consisting of Alpha Modification of Eagle’s Medium (αMEM; Thermo Fisher Scientific, Scoresby, Australia) supplemented with 10% (v/v) foetal bovine serum (FBS; Thermo Fisher), 2 mM L-glutamine (Life Technologies Inc.), 100 U/mL penicillin, and 100 μg/mL streptomycin (Thermo Fisher). Cells were cultured at 37°C in a humidified atmosphere containing 5% CO_2_. NHBC were cryopreserved in liquid nitrogen and thawed and expanded in culture, as required. Cells were passaged up to a maximum of five passages prior to use in experiments. For all assays, cells were seeded at a density of 2 × 10^4^ cells/cm^2^ in 48-well tissue culture plates.

### Human osteocyte-like cultures

NHBC were induced to differentiate into an osteocyte-like phenotype using a previously established protocol.^(15,48,49)^ Briefly, cells were seeded into 48-well plates and maintained under standard growth conditions until confluent. Culture medium was then replaced with osteogenic differentiation medium consisting of αMEM supplemented with 5% (v/v) foetal bovine serum (FBS; Thermo Fisher), 1.8 mM potassium dihydrogen phosphate (Sigma-Aldrich, St. Louis, MO, USA), 100 μM ascorbate-2-phosphate (Sigma-Aldrich), 2 mM L-glutamine (Life Technologies, Inc.), and penicillin/streptomycin (1 U/mL each; Thermo Fisher). Cells were cultured under humidified conditions at 37°C in an atmosphere containing 5% CO_2_ for up to 4 weeks to promote osteocyte-like differentiation. Differentiation medium was replenished twice weekly throughout the culture period.

### Staphylococcus aureus culture

The methicillin-resistant *S. aureus* (MRSA) strain WCH-SK3^(29)^ was cultured under standard aerobic growth conditions. For routine culture, bacteria were grown overnight in nutrient broth (NB) containing 5 g/L NaCl, 3 g/L beef extract, and 10 g/L peptone (Chem-Supply, SA, Australia) at 37°C with agitation at 200 rpm. Where required, bacteria were also cultured on nutrient broth agar (NBA), consisting of NB supplemented with 1.5% (w/v) bacteriological agar (Sigma-Aldrich). For each experiment, bacterial concentrations were estimated using previously generated standard curves correlating optical density (OD_600 nm_ or OD_630 nm_) with colony-forming units (CFU)/mL. Final inoculum concentrations were confirmed by serial dilution and colony counting on NBA plates.

### Osteocyte infection model

Osteocyte infection experiments were performed, essentially as previously described.^(23)^ In brief, bacterial cultures were harvested by centrifugation at 10,000×g for 10 min and resuspended in sterile PBS to achieve the desired bacterial density corresponding to the required multiplicity of infection (MOI) for each experiment. Prior to infection, cell culture medium was removed, and host cells were washed twice with sterile PBS. Cells were then exposed to the bacterial suspension or PBS alone as an uninfected control. Bacteria were incubated with host cells for 2 h at 37°C to allow infection. Cells were then rinsed twice with PBS to remove non-adherent bacteria and then incubated for a further 2 h at 37°C in antibiotic-free medium containing 10 μg/mL lysostaphin (AMBI Products LLC, Lawrence, NY, USA) to eliminate remaining extracellular bacteria. After extracellular bacterial clearance, lysostaphin-containing medium was removed and replaced with fresh osteocyte differentiation medium supplemented with antibiotics. Culture supernatants collected at 24 h post-infection were plated onto agar plates to confirm sterility and successful removal of extracellular bacteria. During longer-term experiments, culture medium was replaced twice weekly throughout the experimental period, as previously described.^(23)^

### Immunophenotypic characterisation by flow cytometry

Immunophenotypic profiling of NHBC was performed by flow cytometry using fluorochrome-conjugated monoclonal antibodies (MAb) directed against human cell surface markers associated with antigen presentation and immune regulation: HLA-ABC (MHC Class I;)-PE (clone W6/32; eBioscience; Cat. 16998385), HLA-DR-PE-Cy7 (clone L243; BD Biosciences; Cat. 335795), CD154 (CD40L)-APC (clone 89-76; BD Biosciences, Cat. 648887), CD80-PE (clone 307; BD Biosciences; Cat. 559370), CD83-FITC (clone HB15e; BD Biosciences; Cat. 556910), CD86-FITC (clone 2331(FUN -1); BD Biosciences; Cat. 557343), CD206-FITC (clone 19.2; BD Pharmingen; Cat. 551135), and CD209-FITC (clone DCN46; BD Pharmingen; Cat. 551264).

For staining, cells were washed with cold culture medium and incubated for 30 min on ice in fluorescence-activated cell sorting (FACS) buffer consisting of cold PBS supplemented with 0.1% bovine serum albumin (BSA). Following incubation, cells were washed twice with FACS buffer and resuspended in Dulbecco’s phosphate-buffered saline (DPBS; Gibco) for acquisition. To assess non-specific binding, matched mouse MAb isotype controls (BD Pharmingen) were included for each fluorochrome. Data acquisition was performed using a BD FACSCanto™ II flow cytometer, and data were analysed using FlowJo software.

### Immunofluorescence analysis

NHBC were cultured in Lab-Tek II chamber slides (Nunc, Thermo Fisher Scientific, NY, USA) for MHC Class II staining, and in black-walled clear-bottom 24-well imaging plates (VisiPlate 24-well, Thermo Fisher Scientific) for HLA-DM analysis. For osteocyte differentiation studies, cells were cultured in osteogenic differentiation medium for up to 4 weeks prior to immunofluorescence evaluation, as described above. After the indicated culture period, cells were incubated with 100 μL LysoTracker Red DND-99 solution (1:500 dilution in growth medium) for 1 h at 37°C in a humidified atmosphere containing 5% CO_2_. Following incubation, cells were washed three times with PBS. Cells were then fixed using 100 μL of chilled acetone:methanol (1:1, v/v) for 90 s, followed by three additional washes with PBS. Non-specific binding sites were blocked with 200 μL of 5% normal goat serum in PBS for 1 h at room temperature. For MHC Class II detection, cells were incubated overnight at 4°C with the anti-human MHC Class II MAb, FMC52^(50)^ (provided by Dr. Peter Macardle, Flinders University). After washing with PBS, cells were incubated for 2 h at room temperature in the dark with FITC-conjugated goat anti-mouse IgG secondary antibody (ab6785, Abcam). The HLA-DM staining, cells were processed under the same conditions, except that Alexa Fluor 594-conjugated goat anti-mouse IgG secondary antibody (ab150116, Abcam) was used. For osteocyte differentiation imaging, filamentous actin was visualised using phalloidin staining and nuclei were counterstained with DAPI-containing fluorescence mounting medium (Sigma, USA). Fluorescence images were acquired using an Olympus FV3000 confocal microscope (Olympus, Tokyo, Japan).

### MHC Class II expression in *S. aureus-*infected osteocyte-lineage cells

To determine the effects of bacterial exposure on MHC Class II expression, undifferentiated NHBC and 4-week differentiated osteocyte-like cells were infected with *S. aureus*, as described above. At 24 h and 48 h post-infection, cells were washed three times with PBS to remove residual medium and non-adherent material. Cells were then stained with HLA-DR-PE-Cy7 MAb (BD Bioscience) for flow cytometry, as described above.

### Isolation of PBMC and CD4⁺ T cells

Peripheral blood was collected into EDTA-coated tubes and peripheral blood mononuclear cells (PBMC) were isolated by density-gradient centrifugation using Lymphoprep™ (Stemcell Technologies, VIC, Australia). Briefly, whole blood was diluted 1:1 with phosphate-buffered saline (PBS), layered over density-gradient medium and centrifuged at 500 × g for 30 min at room temperature without braking. The PBMC layer was collected, washed twice in PBS and resuspended in complete culture medium. Where required, PBMC were cryopreserved in foetal bovine serum (FBS) containing 10% dimethyl sulfoxide (DMSO) and stored in liquid nitrogen until use. CD4⁺ T cells were isolated from PBMC by immunomagnetic negative selection using the EasySep™ Human CD4⁺ T Cell Isolation Kit (Stemcell Technologies) according to the manufacturer’s instructions. Briefly, non-target cells were labelled with antibody isolation cocktails and magnetic RapidSpheres™, followed by magnetic depletion using the EasySep™ separation system. Purified CD4⁺ T-cell populations routinely exceeded 95% purity as determined by flow cytometry.

### CellTrace Violet labelling and autologous co-culture of CD4⁺ T cells

For proliferation assays, purified CD4⁺ T cells were labelled with CellTrace™ Violet (CTV; Thermo Fisher Scientific). Cells were resuspended at 1 × 10⁶ cells mL⁻¹ in pre-warmed PBS containing 5 μM CTV and incubated for 20 min at 37°C protected from light. Staining was quenched by the addition of five volumes of complete medium containing 10% FBS followed by incubation for 5 min at room temperature. Cells were washed twice in complete medium and used 5 × 10^4^ cells/well for subsequent autologous co-culture experiments.

For autologous proliferation assays, CTV-labelled CD4⁺ T cells were co-cultured with four-week differentiated osteocyte-like cells derived from the same donor. Where indicated, osteocyte-like cultures were infected with *S. aureus* prior to co-culture. CD4⁺ T cells cultured alone served as negative controls, while CD3/CD28 activation beads (Gibco, ThermoFisher, Cat.11131D) were used as positive controls. Following 5 days of co-culture, T-cell proliferation was assessed by flow cytometry and analysed using FlowJo (BD Biosciences).

### Western blot analysis of MHC Class II molecules

Western blot analysis was performed to assess expression of MHC Class II based on previously described methods.^(51,52)^ Total protein was extracted from cell cultures using M-PER lysis buffer (Thermo Scientific, USA). Protein concentrations were determined using a bicinchoninic acid (BCA) protein assay kit (Thermo Scientific, USA), according to the manufacturer’s instructions. 10 µg/well of total protein was separated on 10% SDS– polyacrylamide gels (NuPAGE™ Bis-Tris Protein Gels, Thermo Fisher) and transferred onto Immobilon-P membranes using a Trans-Blot Turbo transfer system (Bio-Rad). Membranes were blocked in 1% (w/v) skim milk prepared in Tris-buffered saline containing Tween-20 (TBST) to reduce non-specific binding, followed by overnight incubation at 4°C with mouse anti-human FMC 52. Following primary antibody incubation, membranes were washed and incubated for 2 h with IRDye 800-conjugated goat anti-mouse secondary antibody (LI-COR Biotechnology, NE, USA). Membranes were then washed thoroughly with TBST prior to imaging. Protein bands were detected using an ODYSSEY CLx imaging system (LI-COR). Densitometric quantification of immunoreactive bands was performed using ImageJ software.

### Immunohistochemistry of human bone tissue

*Ex vivo* human bone specimens were fixed, decalcified and sectioned at a thickness of 10 μm. Sections were blocked with 100 μL of 5% goat serum in PBS for 30 min at room temperature to minimise non-specific antibody binding. Sections were then incubated overnight at 4°C with primary antibodies directed against HLA-DR and HLA-ABC. Following primary antibody incubation, sections were washed with PBS and incubated for 30 min at room temperature in the dark with Alexa Fluor 594-conjugated goat anti-mouse IgG (H&L) secondary antibody (Abcam). Slides were subsequently mounted using fluorescence mounting medium containing DAPI (Sigma, USA) for nuclear counterstaining and examined by fluorescence confocal microscopy. Sections of bone were also stained histologically using the RGB trichrome stain, as previously described.^(32,53)^

### Statistical analysis

Statistical analyses were performed using GraphPad Prism version 10 (GraphPad Software, San Diego, CA, USA). Time course data were analysed using one-way analysis of variance (ANOVA). Comparisons between two groups were performed using Student’s t-test. Data are presented as mean ± SEM unless otherwise indicated. Differences were considered statistically significant for *p* < 0.05.

## Results

### Data screening for MHC Class II-associated antigen-processing pathways in *S. aureus*-exposed human osteocytes

To investigate whether human osteocytes express a transcription profile consistent with antigen-processing and antigen-presentation capabilities, we first screened gene microarray datasets from our previous study,^(23)^ comparing human primary osteocytes acutely exposed to *S. aureus* with untreated controls. As shown in **Table 1**, infection was associated with significant regulation of a broad range of key antigen processing and presentation-associated mediators. Importantly, this included upregulation of the MHC Class II transcriptional regulator *CIITA* (2.37-fold, *p* = 0.0163), together with the MHC Class II genes *HLA-DRA* (16.98-fold, *p* = 0.00279), *HLA-DRB1* (1.69-fold, *p* = 0.0465), and *HLA-DPA1* (1.57-fold, *p* = 0.0231).

A coordinated significant increase was also observed in genes encoding components of the antigen-processing and peptide-loading pathway, including *CD74* (2.94-fold), *HLA-DMB* (1.33-fold), cathepsin S/*CTSS* (34.14-fold), cathepsin L/*CTSL* (2.18-fold), *TAP1* (22.01-fold), *TAP2* (9.49-fold), *TAPBP* (3.53-fold), *ERAP1* (1.79-fold), *PSMB8* (4.90-fold), and *PSMB9* (5.80-fold), consistent with activation of the intracellular machinery required for peptide generation, transport, and loading onto MHC Class II molecules.

Expression of co-stimulatory molecules associated with APC–T-cell interactions was also increased, including *CD40*, *CD80*, *CD83*, and *ICAM1* (all *p* < 0.05), whereas *CD86* expression was not significantly altered. Among the cytokine genes associated with T cell activation, *IL6* showed the largest significant increase (82.97-fold), together with significant increases in *IL7* and *IL15*. Together, this analysis provided a strong rationale for the experimental investigation of MHC Class II-associated protein expression and the immunostimulatory properties of human osteocyte-lineage cells.

### Baseline immunophenotypic characterisation of human bone-derived osteoblasts

To examine the baseline immune-related phenotype of human primary osteoblasts (NHBC), cells were analysed by flow cytometry for expression of MHC molecules HLA-DR and co-stimulatory immune-associated surface markers (**Figure 1**). As expected, NHBC showed strong and uniform constitutive cell surface expression of MHC Class I. Unstimulated NHBC contained a distinct HLA-DR-positive subpopulation (**Figure 1B**), with a mean of 10.6% (range, 5.1–14.8%, **Figure 1C**) of NHBC expressing surface HLA-DR under basal culture conditions. Evaluation of co-stimulatory and immune regulatory markers indicated that CD154/CD40L represented the most prominent immune-associated marker detected, with a mean of 15.6% (range 12.0–17.6%) positive cells (**Figure. 1C**). CD83, CD80, CD209, CD206 and CD86 were all expressed but at lower levels (**Figure 1B**).

**Figure 1.**
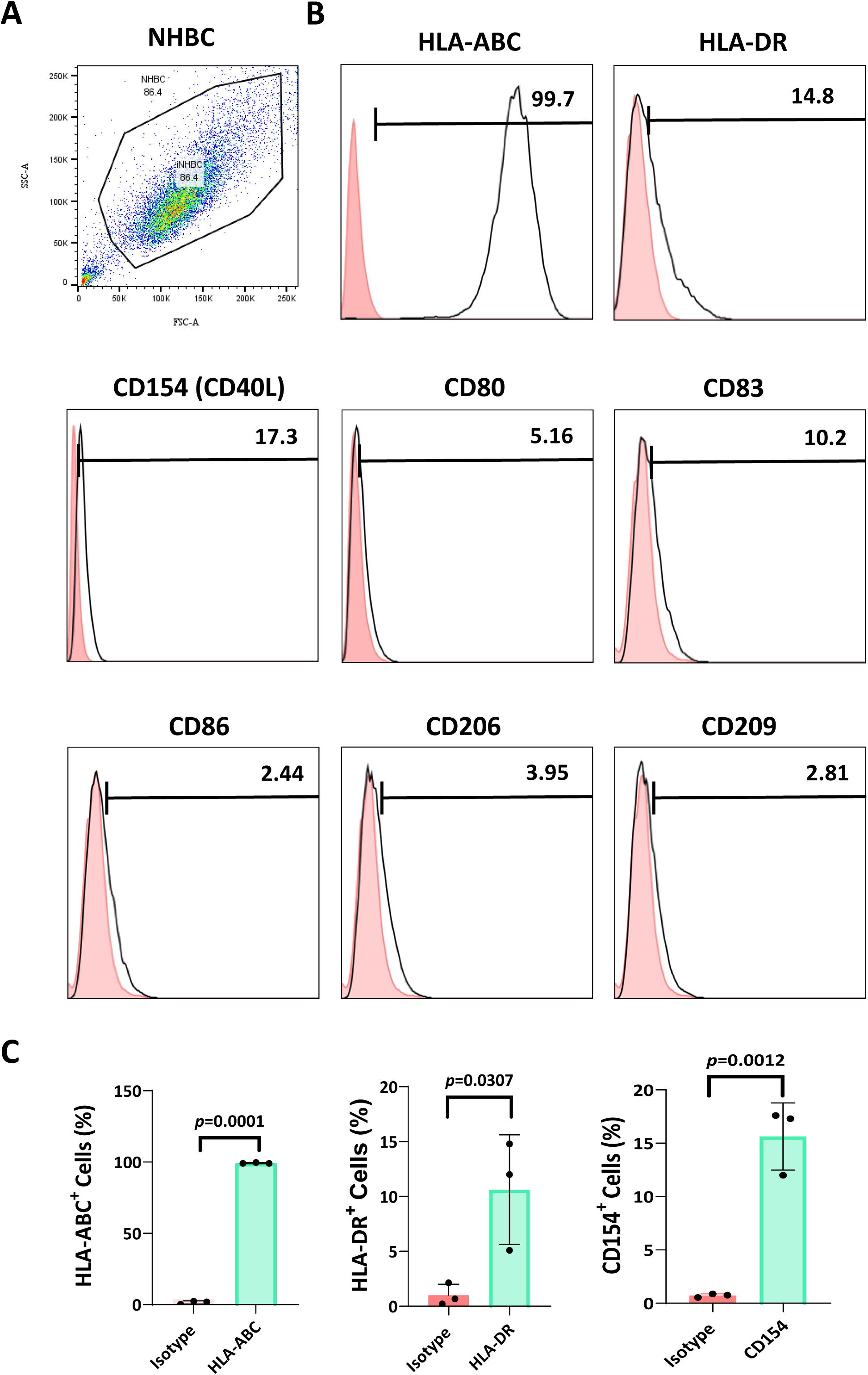
Immunophenotypic characterisation of human primary bone-derived osteoblasts (NHBC). A) Representative flow cytometry gating strategy used to identify the NHBC population for immunophenotypic analysis. B) Representative flow cytometry histograms showing cell surface expression of antigen-presentation and immune-associated markers in NHBC. Black histograms represent specific antibody staining, and red shaded histograms represent matched isotype controls. Expression of HLA-ABC (MHC class I), MHC class II molecule including HLA-DR, and co-stimulatory molecules including CD80, CD83, CD206, CD209 and CD154 (CD40L) is shown. Numbers indicate the percentage of positively stained cells within the gated NHBC population. C) Quantification of HLA-ABC, HLA-DR, and CD154 expression in NHBC. Data are presented as mean ± SEM from independent donor-derived cultures, with individual data points shown. Statistical significance was determined using Student’s t-test. These data demonstrate strong constitutive expression of HLA-ABC together with lower-level expression of HLA-DR and selected immune-associated surface molecules in human osteoblastic cells.

To further investigate the antigen presentation-associated phenotype of NHBC, *in situ* immunofluorescence staining was performed, which demonstrated nearly uniform intracellular expression of MHC Class II under basal culture conditions (**Figure 2A & B**). To determine whether intracellular antigen-processing machinery was also present, NHBC were stained for the peptide-loading chaperone HLA-DM together with LysoTracker to visualise acidic endo-lysosomal compartments associated with antigen processing machinery; confocal microscopy revealed near uniform expression of HLA-DM, substantially co-localised within lysotracker-positive compartments (**Figure 2C & D)**.

**Figure 2.**
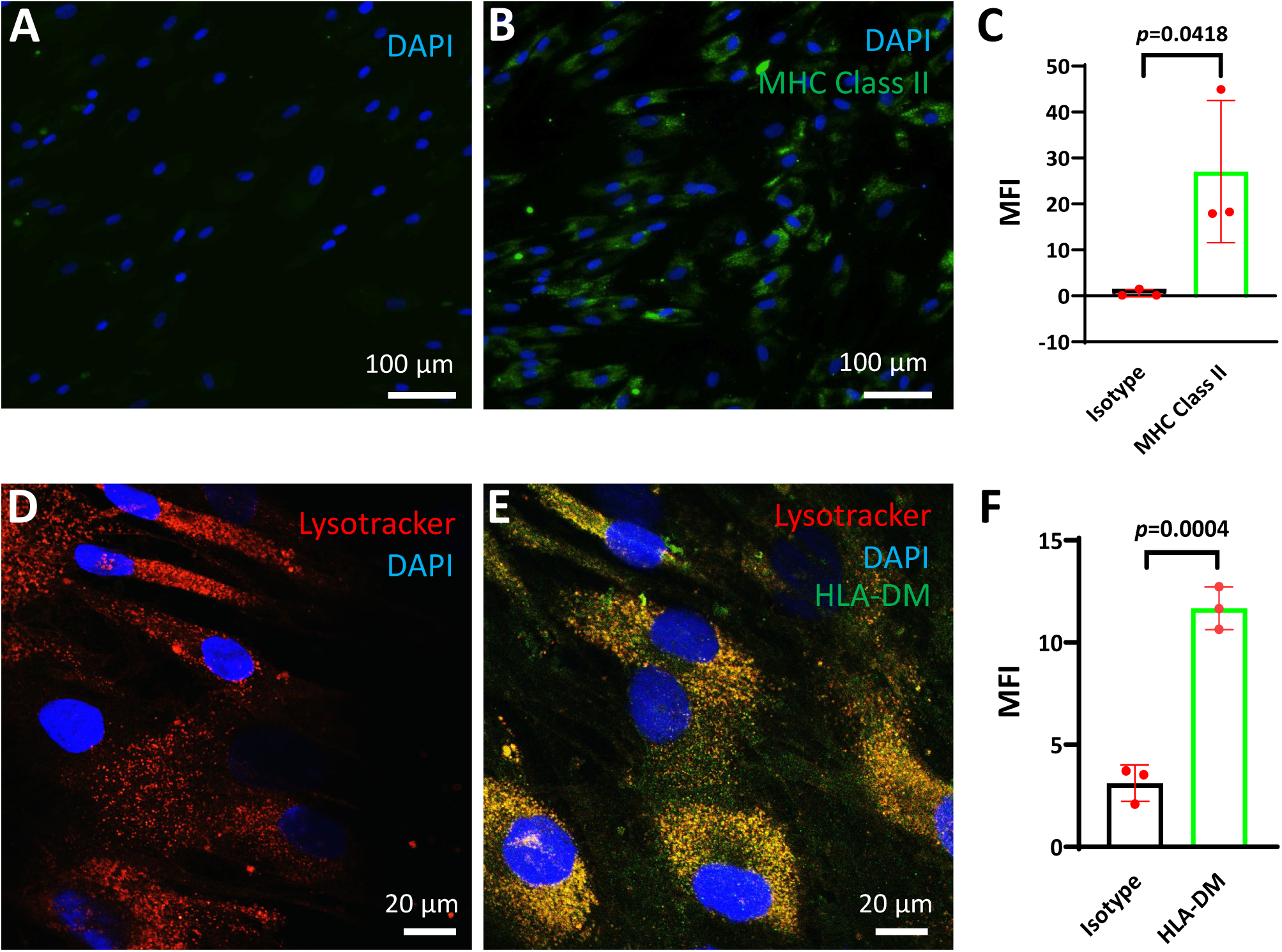
Human primary osteoblasts express MHC class II and HLA-DM under basal conditions. A) Representative immunofluorescence image of NHBC stained with mouse IgG isotype control antibody. B) Representative immunofluorescence image of NHBC stained for MHC class II (green). MHC class II-associated immunoreactivity was detected in a subset of cells under basal culture conditions. Scale bar, 100 μm. C) Quantification of MHC class II-positive NHBC determined by immunofluorescence analysis. D) Representative confocal image of NHBC labelled with LysoTracker (*red*) to identify acidic endosomal/lysosomal compartments while E) confocal image of NHBC stained for HLA-DM (*green*) together with LysoTracker. Merged images demonstrate intracellular HLA-DM localisation within LysoTracker-positive vesicular compartments. Nuclei were counterstained with DAPI (*blue*). Scale bar, 20 μm. F) Quantification of HLA-DM-positive NHBC determined by immunofluorescence analysis. Data are presented as mean ± SEM from independent experiments. Each symbol represents an independent donor culture. Statistical significance was determined using Student’s t-test (*p* < 0.05).

Together, these findings indicate that unstimulated human primary osteoblasts exhibit a restricted but detectable immune-associated phenotype characterised by robust intracellular MHC Class II and HLA-DM expression and limited constitutive cell surface expression of MHC Class II and selected co-stimulatory molecules.

### Mature osteocyte-like cells retain intracellular MHC Class II and HLA-DM-associated machinery

To determine whether differentiation toward an osteocyte-like phenotype influences MHC Class II-pathway expression, protein lysates from NHBC cultured under osteogenic differentiation conditions for up to 4 weeks were analysed by Western blotting. Acquisition of an osteocyte-like phenotype following differentiation was confirmed by increased expression of mature osteocyte marker genes (**Suppl. Figure 1**). A distinct band corresponding to MHC Class II protein was evident in both 2-week and 4-week differentiated cultures, with an apparent but not statistically significant relative increase in the more mature cultures (**Figure 3A & B**), demonstrating that expression is at least maintained with progression to a mature phenotype. Osteocyte-like cultures were further analysed by immunofluorescence confocal microscopy, which showed MHC Class II immunoreactivity distributed within cell bodies and extending along dendritic-like cellular processes, visualised by phalloidin staining (**Figure 3C**). Similarly, HLA-DM staining revealed mature osteocyte-like cell expression, detected as punctate intracellular fluorescence distributed throughout the cytoplasm, with prominent perinuclear localisation and extension into dendritic-like cellular projections (**Figure 3D**).

**Figure 3.**
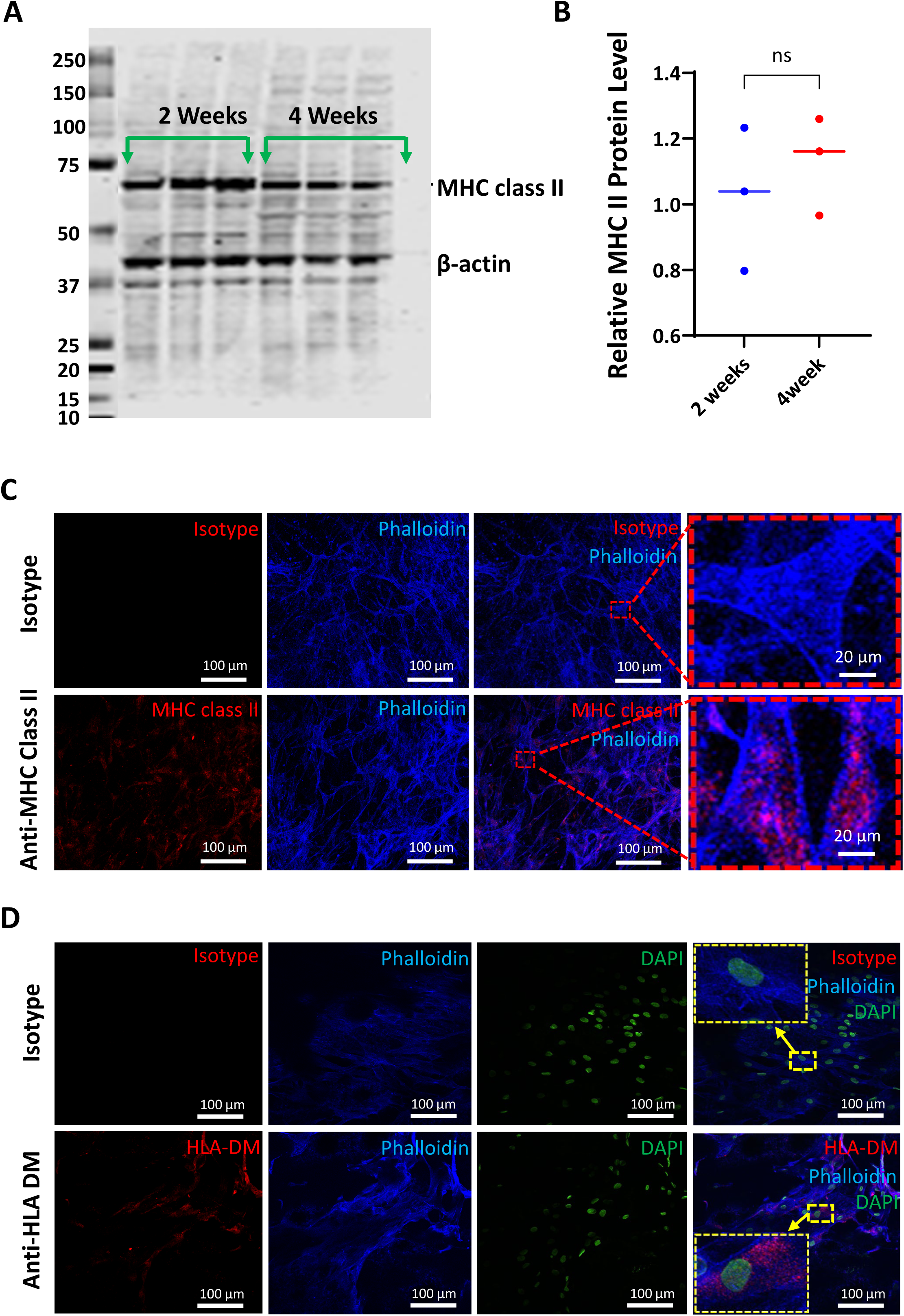
MHC Class II and HLA-DM expression are retained under osteocyte differentiation. A) Representative Western blot analysis of MHC Class II protein expression in NHBC cultures differentiated for 2 and 4 weeks under osteogenic conditions. β-actin served as a loading control. Arrows indicate the respective MHC Class II and β-actin bands. B) Densitometric quantification of MHC Class II protein expression normalised to β-actin. Each symbol represents an independent donor culture, and horizontal bars indicate the mean. Data were analysed using Student’s t-test. (C) Representative confocal immunofluorescence images of 4-week differentiated osteocyte-like cells stained for MHC class II (*red*) and phalloidin-labelled actin cytoskeleton (*blue*). Merged images demonstrate intracellular MHC Class II immunoreactivity throughout osteocyte-like cell bodies and dendritic processes. Enlarged inset panels highlight MHC Class II-associated staining within osteocyte-like cells. Scale bars, 100 μm; inset scale bars, 20 μm. D) Representative confocal immunofluorescence images of 4-week differentiated osteocyte-like cells stained for HLA-DM (*red*), phalloidin (*blue*) and DAPI (*green*). HLA-DM immunoreactivity was detected within osteocyte-like cells and localised throughout the cell body and dendritic processes. Enlarged inset panels highlight intracellular HLA-DM-associated staining. Scale bars, 100 μm. Data are representative of 3 independent experiments performed using cells cultured from different donor bone.

### *S. aureus* exposure increases the presence of MHC class II protein in mature osteocyte-like cells

Having established robust basal expression of MHC Class II and HLA-DM by osteocytes, we next determined whether expression is altered in response to bacterial challenge. Using an established model,^(23)^ osteocyte-like cells were subjected to a one-hour exposure period to *S. aureus,* whereupon extracellular bacteria were removed. MHC Class II protein expression was then assessed after 24h and 48h post-exposure by Western blotting. Significant increases in relative MHC Class II protein levels were observed in *S. aureus*-exposed cultures compared with corresponding non-infected controls, with the strongest increase observed at the later post-infection time point (**Figure 4A & B**).

**Figure 4.**
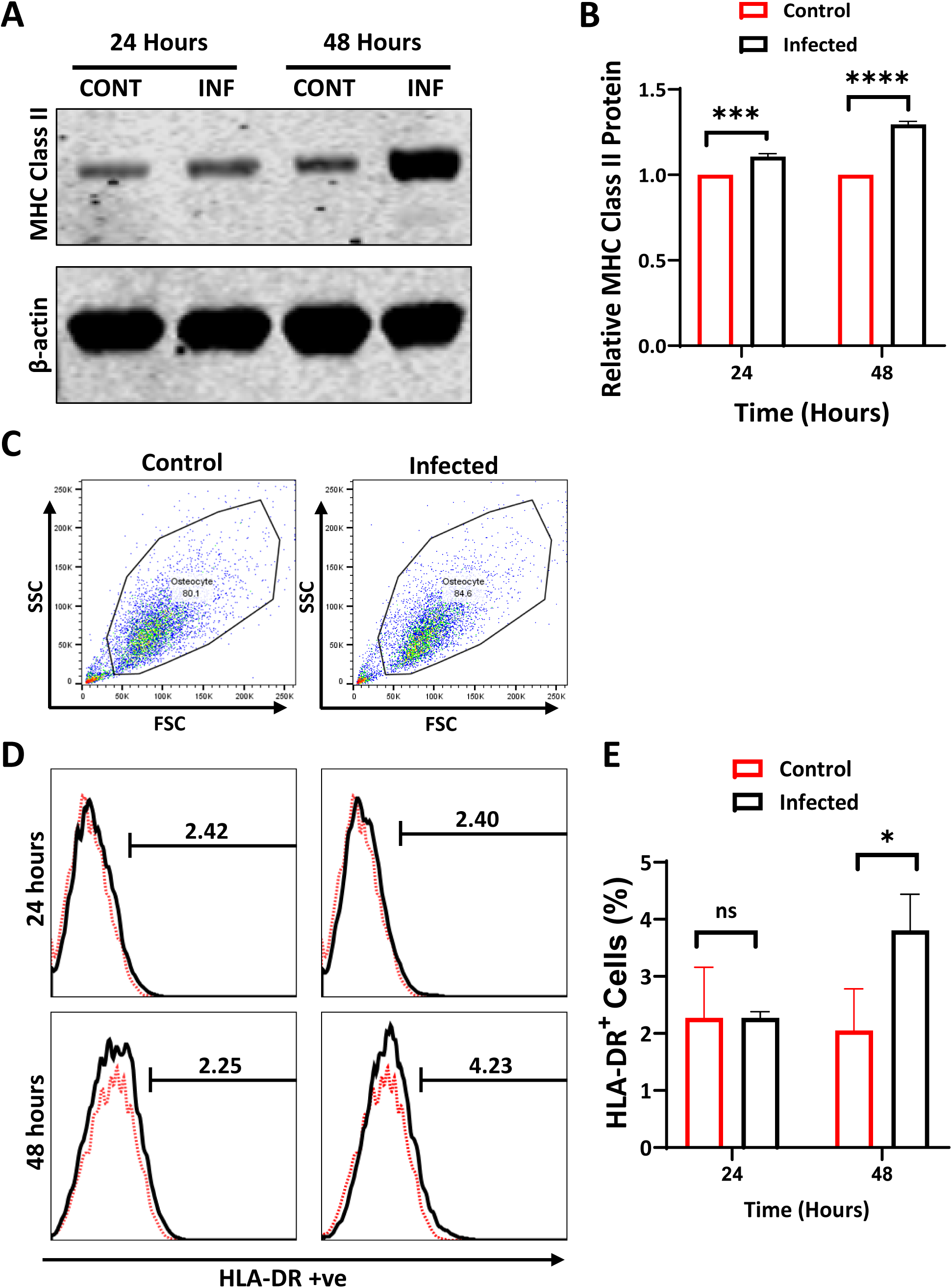
*S. aureus* exposure enhances MHC Class II expression by osteocyte-like cells. A) Representative Western blot analysis of MHC Class II protein expression in four-week differentiated osteocyte-like cells cultured under control (CONT) conditions or following infection with *S. aureus* (INF). β-actin was used as a loading control. B) Densitometric quantification of MHC Class II protein expression was performed normalised to β-actin levels. Data are expressed as means ± SEM relative to control osteocyte-like cells. C) Representative flow cytometry gating strategy showing the osteocyte population under control and infected conditions. D) Representative flow cytometry histograms showing cell surface HLA-DR expression in osteocyte-like cells cultured under control conditions or following *S. aureus* infection. Numbers indicate the percentage of HLA-DR-positive cells within the gated osteocyte population. Red dotted histograms represent isotype controls. E) Quantification of HLA-DR-positive osteocyte-like cells under control and infected conditions. Data are presented as means ± SEM of 3 independent experiments. Statistical significance was determined using Student’s t-test, denoted by \**p* < 0.05; \*\*\**p* < 0.001.

To determine whether this increase was accompanied by altered cell surface expression, the strongly adherent osteocyte-like cells were enzymatically detached and analysed by flow cytometry for surface HLA-DR expression. *S. aureus* exposure did not detectably alter the osteocyte light scatter characteristics, consistent with little effect on cell viability (**Figure C**). The proportion of HLA-DR-positive cells across independent experiments modestly but significantly increased by 48 h in response to bacterial exposure (**Figure 4D & E**).

### *S. aureus*-exposed osteocyte-like cells induce autologous CD4⁺ T-cell proliferation

Co-culture experiments were performed to determine whether osteocyte-like cells could promote autologous CD4⁺ T-cell proliferation following bacterial challenge. CD4⁺ T cells were negatively selected from cryopreserved PBMC isolated at the time of patient joint replacement surgery, resulting in populations of approximately 98% purity (**Figure 5A**). Cultured alone, over a 5-day period, purified cells remained largely undivided, demonstrating minimal spontaneous proliferation (**Figure 5B**). Culture of T cells with *S. aureus* also had no effect on proliferation (**Figure 5C**), confirming that no significant contaminant haemopoietic APC were present in the preparations, whereas CD3/CD28-bead pan-stimulation induced the expected robust proliferation with multiple sequential division peaks (**Figure 5D**). Co-culture with non-*S. aureus* exposed autologous osteocyte-like cells induced only limited CD4⁺ T-cell proliferation, with most cells remaining within the undivided gate and only a small proportion undergoing cell division (**Figure 5E**). In contrast, *S. aureus*-exposed osteocyte-like cells promoted a marked proliferative response, with the emergence of multiple daughter-cell generations (**Figure 5F**), consistent with osteocyte-expressed MHC Class II-restricted bacterial antigen presentation. These data provide functional evidence that osteocyte-lineage cells can participate in adaptive immune activation in the context of bacterial challenge.

**Figure 5.**
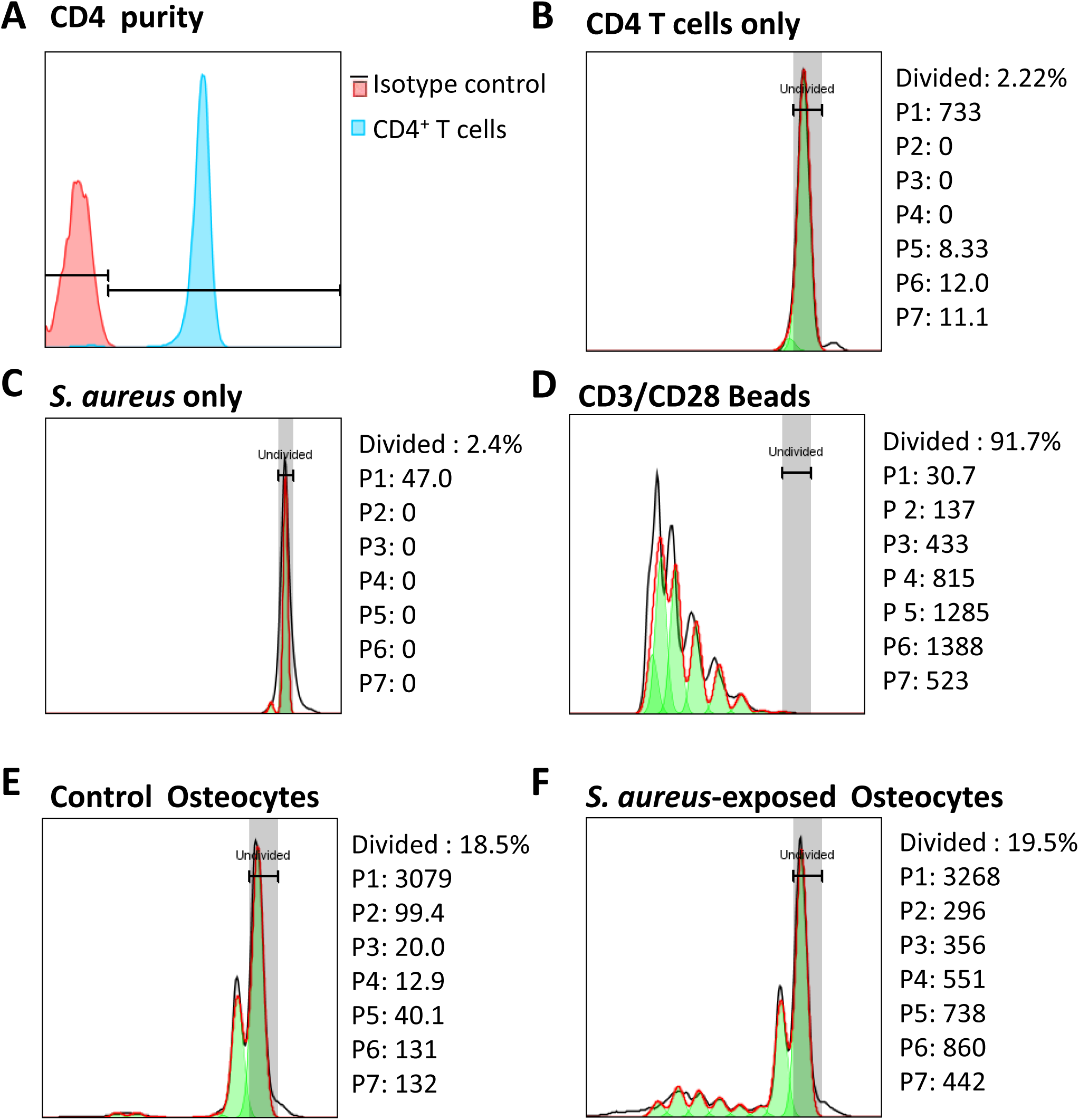
Effect of *S. aureus* exposure of osteocyte-like cells on autologous CD4⁺ T-cell proliferation. Representative CellTrace™ Violet (CTV) proliferation profiles of autologous CD4⁺ T cells following 5 days co-culture with four-week differentiated osteocyte-like cells. CD4⁺ T cells were isolated from the same donor used to generate osteocyte-like cultures and labelled with CTV prior to co-culture. A) Flow cytometry assessment of CD4^+^ T cell purity. B) CD4⁺ T cells cultured alone alone remained largely undivided; C) *S. aureus-*pre-exposed well control (; D) CD4⁺ T cells stimulated with CD3/CD28 pan-activation beads (positive control) exhibited extensive proliferation characterised by progressive CTV dilution and multiple daughter-cell populations; E) Co-culture with uninfected autologous osteocyte-like cells induced limited CD4⁺ T-cell proliferation; (F) Co-culture with *S. aureus*-exposed autologous osteocyte-like cells resulted in increased CD4⁺ T-cell proliferation, evidenced by enhanced CTV dilution and the appearance of multiple proliferative peaks. Histograms are representative of three independent experiments. Grey bands represent the corresponding undivided reference population. Data were acquired by flow cytometry and analysed using FlowJo software. Green shaded peaks (P1–P7) indicate successive generations of divided CD4⁺ T cells.

### Human PJI bone tissue demonstrates osteocyte-associated HLA expression

To determine whether osteocyte HLA-associated immunoreactivity is present in clinically relevant human bone tissue, bone specimens from a PJI patient undergoing revision arthroplasty were obtained proximal to the visually infected site. RGB trichrome staining was performed, a method we recently described as potentially diagnostic of PJI.^(32)^ RGB-stained bone exhibited large swathes of degraded collagen matrix (*red* stain) between lamellae, as well as distinct perilacunar ‘collars’ of degraded matrix with rounded osteocyte lacunae, however with preservation of osteocyte-occupied lacunae throughout the matrix (**Figure 6A**). Sections were then immunostained with isotype controls, anti-MHC Class I (HLA-ABC) and anti-MHC Class II (HLA-DR) (**Figure 6B-D**). As expected, extensive HLA-ABC immunoreactivity was detected in regions containing multiple osteocyte lacunae, pervading the bone matrix and consistent with canalicular staining (**Figure 6C**). HLA-DR staining was also detected but within a subset of osteocyte lacunae and tended to be in regions close to inflammatory cellular infiltrates (**Figure 6D**). These findings provide clinical evidence that osteocytes embedded within infected human bone tissue express MHC Class II, supporting osteocytes as active participants in adaptive immune responses during bone infection.

**Figure 6.**
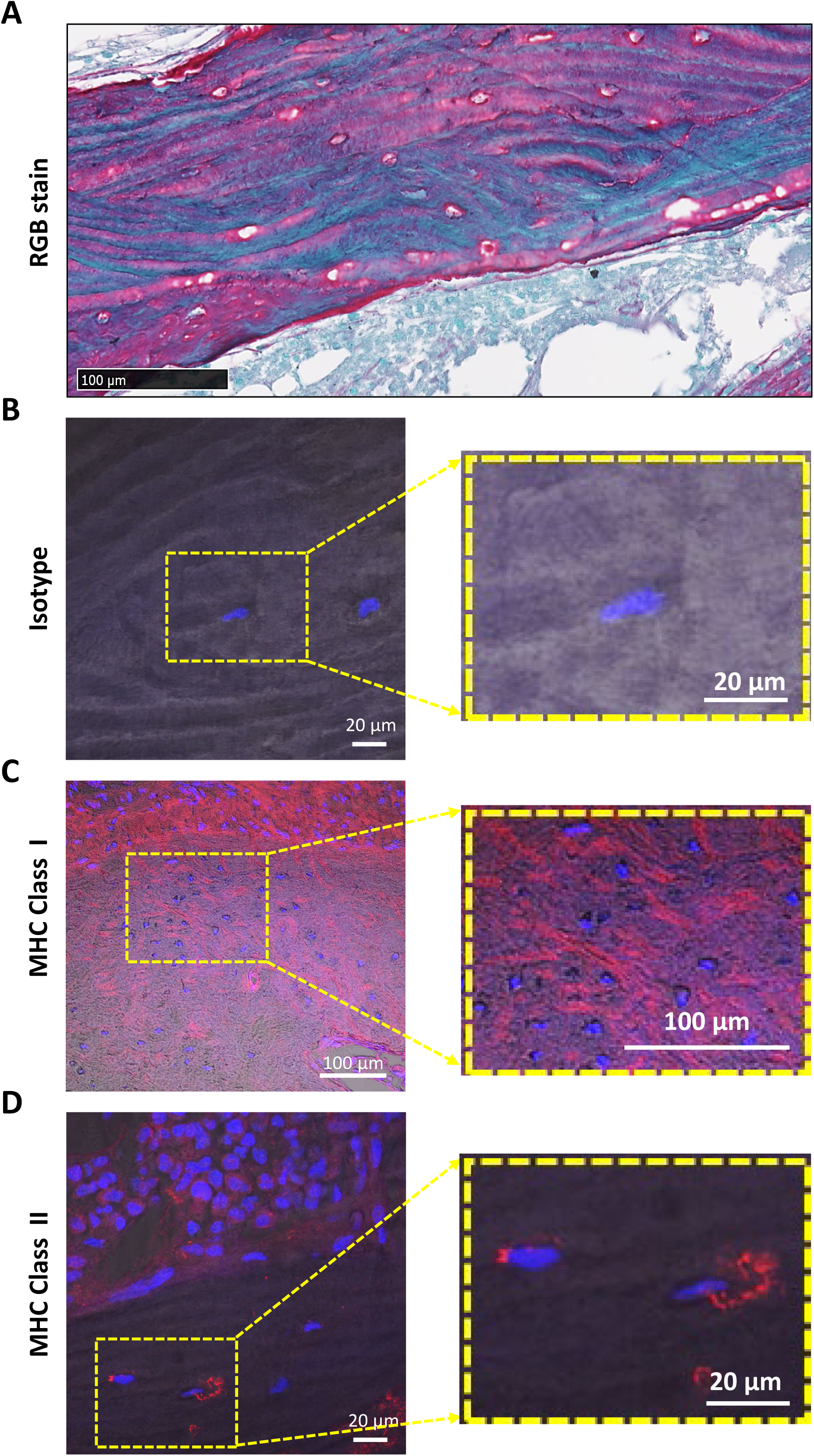
HLA-associated immunoreactivity is detected within osteocyte-containing regions of human prosthetic joint infection (PJI) bone. A) Representative RGB trichrome staining of bone tissue obtained from a PJI patient, demonstrating regions of degraded bone collagen matrix (*red* stain) with partial preservation of mature, intact bone matrix (*blue-green* stain), with osteocyte-containing lacunae bordered by thick degraded matrix collars. An inflamed bone marrow is evident. Scale bar 100 μm. B) Representative confocal immunofluorescence image of a serial bone section stained with mouse IgG isotype control antibody. C) Representative image of a serial bone section showing widespread immunostaining for HLA-ABC (MHC class I). D) Representative confocal image of a serial bone section stained for HLA-DR (MHC class II). HLA-DR-associated immunoreactivity was observed adjacent to osteocyte lacunae within infected bone tissue. Inset images are enlarged as indicated. Nuclei were counterstained with DAPI (blue). Scale bars are indicated for each image.

## Discussion

The present study provides evidence that human osteocyte-lineage cells possess previously unrecognised immunophenotypic properties including basal and inducible expression of MHC Class II, antigen processing and T-cell costimulatory pathways. Analysis of published gene expression datasets revealed that human bone explant-derived primary osteocytes, exposed acutely to *S. aureus*,^(23)^ differentially regulated the expression of a wealth of genes associated with antigen processing and MHC Class II-restricted presentation. This included at least five MHC Class II genes, genes involved in lysosomal trafficking, peptide editing and loading, and T-cell co-stimulation, proliferation and activation. We confirmed that human primary osteoblast-lineage cells express basal MHC Class II-associated features, and importantly, that these features are maintained following differentiation to a mature osteocyte stage. Critically, bacterially-exposed mature osteocyte-stage cells could promote autologous CD4⁺ T-cell proliferation. Osteocytes in human PJI bone were shown also to express MHC Cass II, strongly suggesting that these processes occur *in vivo*.

MHC Class II expression has been reported in several osteoblast-lineage^(8)^ and non-professional antigen-presenting stromal cell populations,^(39,40,54)^ and this may permit rapid responses to pathogens in the bone marrow. Previous studies have established that mesenchymal and osteoblast-lineage cells can acquire components of an inducible MHC Class II antigen-processing programme, including HLA-DR/MHC Class II, the CD74 invariant chain, CLIP-dependent peptide-loading steps and, in the context of inflammatory cytokine or microbial stimulation, functional T-cell stimulatory capacity.^(8,55)^ Recent work in human adipose-derived MSCs further showed that extracellular matrix cues can promote HLA-DR and CD74 expression, supporting the concept that stromal antigen-presenting phenotypes are shaped by microenvironmental context.^(56,57)^ In line with this broader literature, our study confirms that primary human osteoblast-lineage cells express MHC Class II-associated proteins and extends these observations by demonstrating intracellular HLA-DM localisation within LysoTracker-positive acidic vesicular compartments. HLA-DM is critical for antigenic peptide exchange and stabilisation of MHC Class II-peptide complexes in late endosomal/lysosomal compartments.^(35,55,56,58)^ Our findings therefore support the presence of HLA-DM-associated peptide-editing component of the MHC Class II pathway in human osteoblast-lineage cells.

Importantly, intracellular MHC Class II and HLA-DM-associated staining were retained following differentiation into mature osteocyte-like cells. These observations indicate that antigen-processing machinery and associated intracellular immune-associated pathways are preserved in osteocytes. Localisation of both MHC Class II and HLA-DM throughout the osteocyte cell body and dendritic-like processes further suggests that osteocytes maintain widespread intracellular immunophenotypic organisation. These findings are important because mature osteocytes are by far the most abundant bone cell type, are long-lived and embedded throughout the bone matrix. Bone is also relatively poorly vascularised yet susceptible to bacterial penetrance during infection,^(59)^ and osteocytes therefore likely represent a vital population for ongoing local immune surveillance.

Consistent with our previous investigation,^(23)^ exposure to *S. aureus* induced clear upregulation of MHC Class II/HLA-DR protein expression in mature osteocyte-like cultures. This inducible response is highly relevant to osteomyelitis and PJI, where *S. aureus* is a dominant pathogen and where bacteria may persist within bone despite immune activation and antimicrobial treatment. The relatively small but significant increase in the percentage of osteocytes positive for cell surface expression of HLA-DR following exposure to *S. aureus* compared with total MHC class II protein abundance may reflect intracellular retention, delayed trafficking, or possibly, technical loss of dendritic cellular processes during cell detachment for flow cytometry analysis.

Interestingly, analysis of gene datasets revealed that human osteocytes also express *CTSS* encoding cathepsin S, a key protease involved in the degradation of CD74 and the generation of CLIP-containing MHC Class II intermediates, and an important component of the antigen-processing machinery in professional antigen-presenting cells. Remarkably, *CTSS* gene expression increased approximately 34-fold following *S. aureus* infection. This finding is particularly relevant when considered alongside the localisation of HLA-DM within acidic intracellular compartments and the induction of HLA-DR expression following bacterial exposure. Critically, *S. aureus*-exposed osteocytes induced selective proliferation of autologous CD4⁺ T cells, whereas exposure of CD4⁺ cells to *S. aureus* alone did not induce proliferation, arguing against their non-specific activation by potential *S. aureus* superantigens or the presence of contaminating haemopoietic APC as explanations for the observed response. Together, these observations form a compelling and biologically coherent picture: osteocytes possess key components of the MHC Class II antigen-processing pathway and, following exposure to *S. aureus*, appear capable of processing bacterial antigens and stimulating a selective CD4⁺ T-cell response.

Supportive of the physiological relevance of our *in vitro* findings, HLA-DR expression was also clear in osteocytes *in situ* in PJI patient bone. This observation indicates that osteocytes can express or acquire adaptive immunity-associated phenotypes *in vivo*. In this context, our group has shown that in human PJI, bone exhibits specific osteocyte-associated bone matrix degradation and peri-lacunocanalicular remodelling responses, mediated in part by host matrix metalloprotease (MMP) release, a response we postulated may promote bone fluid flow and allow the influx of immune cells.^(31,32)^ The precise relationship between the observed bone matrix and osteocyte morphological changes in PJI, which seem to be widespread, and the osteocyte expression of MHC Class II, and by inference antigen presentation, which appears more restricted, remains to be established but will be important for the complete respective elucidation of these responses.

Persistent rather than transient induction of MHC Class II expression may be important in the context of chronic bone infection, where *S. aureus*, as well as a number of other bacterial pathogens, can survive intracellularly and evade host clearance.^(22,23,27,28,30)^ In this setting, osteocytes could contribute to ongoing antigen persistence and inflammatory signalling, together with recruitment of immune cells.^(23)^ Furthermore, as the ratio of osteoblasts to osteocytes decreases with advanced age,^(60)^ osteocyte expression of APC machinery may become increasingly important in local bone immune defence throughout our lifetime.

Several limitations of this study should be acknowledged. Firstly, the *in vitro* cell models utilised cannot fully recapitulate the complexity of the osteocyte lacunocanalicular niche *in vivo*, however the use of validated human primary cell models and the observation of clear MHC Class II immunostaining of osteocytes in human clinical bone somewhat mitigate this limitation. Secondly, the pathway(s) of antigen processing in the *S. aureus*-exposed osteocytes was not determined. We have previously demonstrated *S. aureus* internalisation by similar osteocyte cultures to those utilised here.^(23)^ We have shown the involvement of autophagy in the response of host cells but this could not fully explain the fate of internalised bacteria in terms of number or viability; modulation of autophagy had some effect on bacterial escape in long-term infection experiments.^(27)^ *S. aureus* is the predominant pathogen in adult human osteomyelitis and PJI, and its prevalence is likely contributed by its ability to evade host immune defences, and osteocytes appear to be no exception. Therefore, studies with lower virulence pathogens might shed further light on the effectivity of osteocyte-mediated immunity. Relatedly, beyond CD4 T cell proliferation, it will be important to determine the nature of the ensuing immune response, and whether osteocyte presentation of antigen results in disease resolution or is tolerogenic, contributing to a chronic disease state.

## Conclusions

Our findings identify human osteocytes as previously unknown adaptive immune-responsive cells capable of activating inducible MHC Class II-associated pathways during bacterial challenge. Osteocytes were found to exhibit a comprehensive network of gene expression related directly to such a function and were confirmed here to express basal cell surface and cytoplasmic HLA-DR expression, intracellular HLA-DM-linked antigen-processing machinery, and the ability to stimulate CD4-positive T-cell proliferation in response to *S. aureus* exposure. The detection of osteocyte MHC Class II in human PJI bone further supports the clinical relevance of these findings. Given their abundance, longevity and extensive distribution throughout mineralised tissue, osteocytes appear to represent a previously unrecognised immune-responsive cellular network within bone. Together, these findings significantly extend the current understanding of the roles of osteoblast lineage cells in general, and osteocytes in particular, beyond skeletal homeostasis and support a key function of these cells in both innate and adaptive immune responses. These findings expand current concepts of osteoimmunology and establish a foundation for future studies investigating osteocyte-mediated regulation of adaptive immune responses during chronic skeletal infection and inflammatory bone disease.

## Acknowledgements

The authors thank Dr Patrick Asare (Adelaide University) for providing reagents for Western blot analysis, Dr. Randall Grose (Adelaide University and the South Australian Health and Medical Research Institute) for assistance with flow cytometry, and nursing staff of the Department of Orthopaedics and Trauma, Royal Adelaide Hospital for help with participant recruitment and specimen collection. M.A.H. and Q.S. were supported by University of Adelaide Postgraduate Research Scholarships and National Health and Medical Research Council-funded top-up scholarships.

## Author Contributions

M.A.H., P.R.H. and G.J.A. contributed to study conception and design. M.A.H. performed the experiments, data analysis and prepared the initial draft of the manuscript. D. M. and Q. S. performed histology. B.R. and L.B.S. provided patient materials. P.R.H. co-supervised the study, and contributed to primary data analysis. G.J.A. co-supervised the study and is the communicating author. P.R.H. and G.J.A. contributed to manuscript development and all authors contributed to manuscript editing.

## Data availability

The datasets generated and/or analysed during the current study are available from the corresponding author upon reasonable request.

---

**Supplementary Figure S1.**
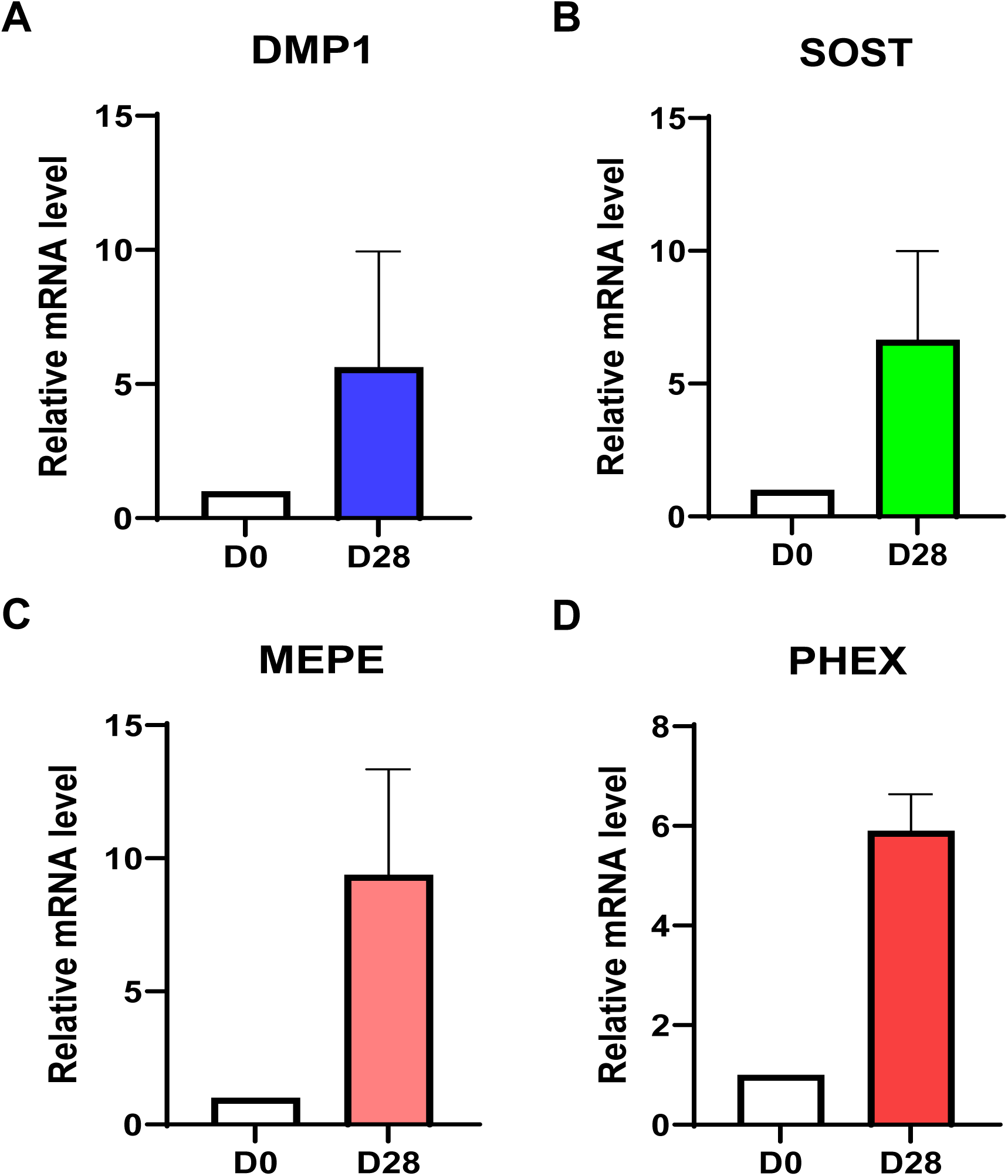
Expression of mature osteocyte markers following in vitro osteogenic differentiation. Primary human bone-derived osteoblasts (NHBC) were cultured under osteogenic differentiation conditions for 28 days to generate osteocyte-like cells, as described in Material and Methods. Total RNA isolation, reverse transcription and quantitative real-time PCR (qRT-PCR) was performed, as described. Fold-change in gene expression in osteocyte-like cultures (D28) relative to undifferentiated NHBC (D0) was calculated using the 2−ΔΔCt method for A) *DMP1*, B) *SOST*, C) *MEPE* and D) *PHEX*. Data are presented as mean values from 3 independent donor-derived cultures. These data confirm acquisition of an osteocyte-like phenotype during differentiation.

